# Bee and Flowering Plant Communities in a Riparian Corridor of the Lower Rio Grande River (Texas, USA)

**DOI:** 10.1101/2020.01.04.894600

**Authors:** Amede Rubio, Karen Wright, Scott Longing

**Affiliations:** Texas A&M International University., Laredo, TX; Texas A&M University, College Station, TX; Texas Tech University, Lubbock, TX

**Keywords:** Lower Rio Grande, bee communities, riparian and upland habitats, diversity

## Abstract

The Rio Grande in Texas serves as the geo-political boundary between the United States and Mexico. It is considered one of the world’s most at-risk rivers and has been the subject of intensified management by the inhabitants of both countries lining its banks. Additionally, invasion by non-native *Arundo donax* (Linnaeus) (Cyperales: Poaceae), giant reed, has been extensive in the riparian corridor, with potential impacts to native wildlife. Locally, there remains a significant lack of ecological community data of riparian and upland habitats parallel to the river. We sampled bee and flowering plant communities monthly over two years, along a 3.22 km stretch of the lower Rio Grande in Webb County, TX. Data show that bee and plant richness and abundance was highest during March-April and September among both habitat types. Analysis of bee communities showed low spatial and temporal variation at the habitat level. Although common bee taxa (Halictidae and Apidae) were numerically dominant, NMS and ISA found key bee species driving community patterns. This included higher abundances of two species in the riparian habitat *Anthophora occidentalis* (Cresson) (Hymenoptera: Apidae) and *Lasioglossum sp.L* (Curtis) (Hymenoptera: Apidae) and one showing affinity for the upland habitat *Halictus ligatus* (Say) (Hymenoptera: Halictidae). Additionally, ISA analysis of plant data revealed that three species were significant indicator taxa in riparian habitats. Further analysis showed a positive correlation between bee generic richness and abundance with various climate attributes. Management of the riparian corridor and associated watershed could include significant areas for ecological restoration to assist pollinators.

## Introduction

Flowering plants and their associated pollinators are intricately linked by evolved mutualisms (Potts et al. 2010, Fiedler, Landis, and Arduser 2012). Pollination is a vital ecosystem service provided by bees that sustains plant communities and contributes to the production of many agricultural crops (Kremen, Williams, and Thorp 2002). This pollinator-plant interdependence may directly influence seed production and genetic variation within managed and wild plant communities (Kremen et al. 2002). It is estimated that bees pollinate over half of the world’s crop varieties and are responsible for an estimated 15 billion dollars in annual revenue (Kremen, Williams, and Thorp 2002, Losey and Vaughan 2006, Kimoto et al. 2012). In addition to managed systems, pollination of wild flowering plant communities are especially dependent on bees (Kremen et al. 2002). Currently, global threats to pollinators are expected to continue if current environmental trends go unmitigated (Potts et al. 2010), which will lead to the reduction of valued ecosystem services provided by bee and other insects (Losey and Vaughan 2006).

The European honeybee, *Apis mellifera* (Linnaeus) (Hymenoptera: Apidae), has received attention because managed colonies in the United States have shown winter declines of over 50% in recent years (Ragsdale, Hackett, and Kaplan 2007). Concurrent with reported losses of honeybee colonies, several native bee species have been listed as targets for conservation due to severely reduced population ranges and sizes (Cameron et al. 2011). Agricultural intensification coupled with increased pesticide use have become rising threats to native bees due to their non-target effects (Hladik, Vandever, and Smalling 2016, Begosh et al. 2020, Longing et al. 2020). Moreover, managed bees can affect wild native bees through vector disease causing agents during foraging and contact with shared floral resources (Fürst et al. 2014). Concomitantly, rampant habitat fragmentation and invasion by non-native plant species will only intensify the decline of native bee populations (Potts et al. 2010). Further losses of pollinators could dramatically affect ecosystem function, therefore understanding wild bee communities is an important area of research to support both the conservation of biodiversity and ecosystem requirements.

The Rio Grande begins in the San Juan Mountains of Colorado and travels approximately 3,200 kilometers before draining into the Gulf of Mexico. It serves as the geographical and political boundary between the United States and Mexico (Karatayev, Miller, and Burlakova 2012). The river and its associated riparian corridors are some of the most anthropogenically influenced and understudied systems in the world (Karatayev, Miller, and Burlakova 2012). The river is also a primary source of drinking water and supports much of the municipal, industrial, and agricultural water needs for major cities along the U.S.-Mexico border. The Rio Grande and associated riparian ecosystems provide resources to maintain food webs, create refugia and habitat for animals, and serve as a steady source of available freshwater (Ellis, Crawford, and Molles Jr 2001). Over-extraction of freshwater, pollution, invasive plant species and climate change continue to influence the Rio Grande (Karatayev et al. 2012, Wilson, Addo-Mensah, and Mendez 2015). Although impacts from anthropogenic activities are widespread, the riparian corridor of the Rio Grande remains understudied regarding its flora and fauna. A need exists to understand how the riparian corridor supports resources for wildlife.

The Rio Grande corridor can be sub-divided into riparian and upland habitats, that are generally distinguished by relative distances to the riverbank composition of the plant community. The vegetation of the Rio Grande riparian habitat consists in part of the following species: woody species sugar hackberry (*Celtis laevigata* Willdenow) (Urticales: Ulmaceae), retama (*Parkinsonia aculeata* Linnaeus) (Fabales: Fabaceae), Mexican ash (*Fraxinus berlandieriana* de Candolle) (Scrophulariales: Oleaceae), black willow (*Salix nigra* Marshall) (Salicales: Salicaceae), granjeno (*Celtis pallida* Torrey) (Urticales: Ulmaceae), and forb species pigeon berry (*Rivina humilis* Linnaeus) (Caryophyllales: Phytolaccaceae), narrowleaf globe mallow (*Sphaeralcea angustifolia* Cavanilles) (Malvales: Malvaceae), common sunflower (*Helianthus annuus* Linnaeus) (Asterales: Asteraceae) (Woodin, Skoruppa, and Hickman 2000, Everitt, Drawe, and Lonard 2002, Racelis et al. 2012). The upland habitat is generally higher in elevation and often is the outermost boundary of the riparian corridor. Upland vegetation consists of woody species, dominated by sugar hackberry (*Celtis laevigata* Willdenow) (Urticales: Ulmaceae), honey mesquite (*Prosopis glandulosa* Torrey) (Fabales: Fabaceae), black brush acacia (*Vachellia rigidula* Bentham) (Fabales: Fabaceae), and forb species such as mock vervain (*Glandularia quadrangulate* Heller) (Lamiales: Verbenaceae), annual sowthistle (*Sonchus oleraceus* Linnaeus) (Asterales: Asteraceae) and plains lazy daisy (*Aphanostephus ramosissimus* de Candolle) (Asterales: Asteraceae) (Everitt, Drawe, and Lonard 1999, Everitt, Lonard, and Little 2007, Racelis et al. 2012). Many plants within the riparian and upland habitats likely provide resources to pollinators, but this has not been determined in our study region.

Grasses in the riparian corridor are dominated by the invasive giant reed (*Arundo donax* Linnaeus) (Cyperales: Poaceae) and guineagrass (*Urochloa maxima* Jacquin) (Cyperales: Poaceae) (Everitt et al. 2011). *A. donax* is widely distributed and has significantly fragmented the landscape by producing large expanses of monotypic stands. Furthermore, invasive buffelgrass (*Cenchrus ciliaris* Linnaeus) (Cyperales: Poaceae) and bermudagrass (*Cynodon dactylon* Linnaeus) (Cyperales: Poaceae) are frequently encountered where rare patches of (non-Arundo) vegetation occurs. Studies have suggested that the rapid growth and spread of invasive grasses can have a severe negative impact on floral resources for pollinators (Fierke and Kauffman 2006), including those along the Lower Rio Grande (LRG) in Texas (Rubio et al. 2014). Due to the pervasiveness of giant reed, *A. donax*, in the riparian corridor, this study aimed to document bee and flowering plant communities over time and in relation to different habitats generally defined by soil and vegetation.

Currently, little is known about the present state of flowering plant and bee communities provided by the riparian corridor, and information produced from related studies could support strategies for restoring disturbed soil and development of new infrastructure. The objectives of this study were to survey bees and plants in the riparian and upland habitats to document spatial and seasonal community differences. Insect pollinators and flowering vegetation are potential biological targets in this intensively managed system, and findings would support conservation through strategies for ecological restoration.

## Materials and Methods

### Study Area

This study was conducted along a 3.22 km stretch of the Lower Rio Grande river (LRG) in southwestern Webb County, TX (27.5013°N; 099.52697°W). The area is situated within the City of Laredo, TX and in proximity to Laredo College campus. The climate of the region is semi-arid subtropical, with hot summers and mild winters (NRCS 2006). The average annual temperature is 30.2°C and average annual precipitation is 54.7 cm (NRCS 2006). Typically, May and September are the wettest months, averaging 7.26 cm of precipitation combined (NRCS 2006). The LRG soil series primarily dominates both habitats, which is described as deep, well drained, very fine sandy loam, and moderately alkaline (Sanders and Gabriel 1985). The LRG’s soil can sustain riparian vegetation through periods of prolonged drought due to its flood water holding capacity (Moore et al. 2016). Gravel has been introduced into the area for the construction and maintenance of access roads by the Department of Homeland Security. The unique LRG plant communities provide suitable habitat for many vertebrate and invertebrate species, including polyphagous beetles (Osbrink et al. 2018) and native bee communities (Henne, Rodriguez, and Adamczyk 2012). The total area representative of the sampled habitats was approximately 3000 m^2^.

Both riparian and upland habitats designated in this study occur within the broader riparian corridor of this river, yet differences in elevation, percent sandy substrate, and vegetation supported our stratification of habitats. Upland habitat ranged between 180 and 530 m from the main stem of the river, while riparian zone habitats were located 50 to 130 m from the river. Sampling plots between each habitat were separated on average by 172 m along linear transects generally running parallel with the river.

### Field Methods

In February 2017, twenty (10 in each habitat) 50 m-long x 1m-wide belt transects were established parallel to the Rio Grande in riparian and upland habitats, to in order to sample monthly at these locations the flowering plant and bee communities. In March – May 2017 24 triplet pan (“bee bowl”) trap stations (12 in riparian zone and 12 in upland terrace zone) were placed 50 m apart within the sampling area (Fig.1). Pan traps were deployed monthly between February 2017 to May 2019 at these locations.

**Fig. 1.**
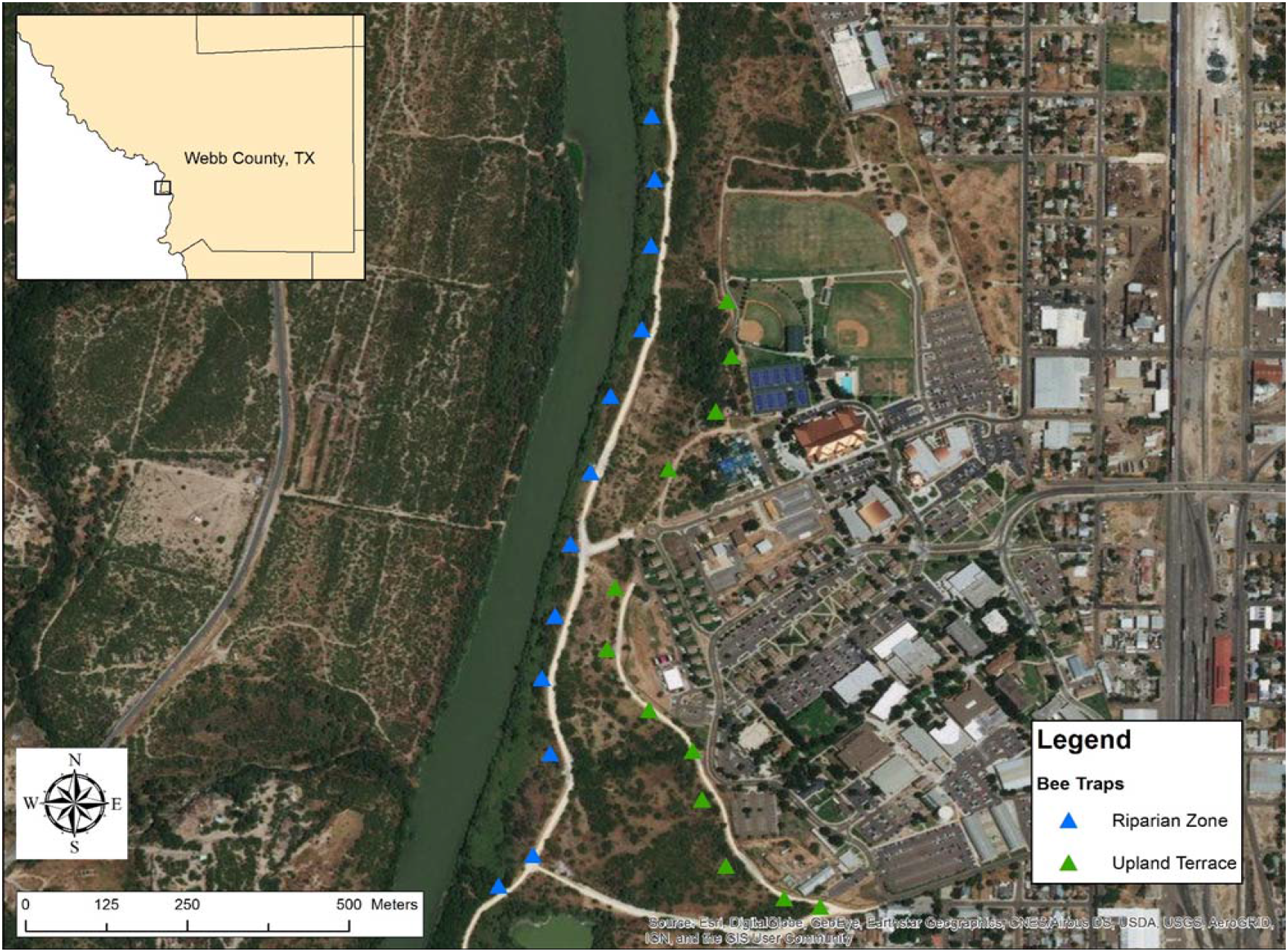
Map of the study area with bee bowl trap locations in riparian and upland habitats (triangles). Locations of vegetation transects for flowering plant surveys (not shown) are within the extent of this sampling area and included habitats within 5 m surrounding bee bowl locations

Vegetation was sampled monthly by walking the length of the belt transects and documenting the presence of plant species in bloom. In addition to transects, blooming flowering plants were also identified in situ around a 5 m radius of each bee pan trap cluster. At pan trap cluster locations, during each visit to collect bee bowl contents (25 total visits), visible blooms were recorded as present, representing one individual count of that plant species. Plants unidentifiable in the field were photographed and/or collected for identification in the lab. Voucher specimens and digital images of flowering plants are being held in the laboratory of Amede Rubio at Texas A&M International University (Dept of Biology and Chemistry).

Bee communities were sampled using aerial nets and bee bowls (i.e. pan traps) (LeBuhn et al. 2016) monthly over two years. Hunt sampling the length of established belt transects was conducted in pairs, one person netting directly from plants and the other recording. Sampling within individual transects lasted approximately 25 minutes. Bee bowls were an adaptation of Droege et al. (2010). Bee bowls are 3.5 oz cups painted three different fluorescent colors (blue, white, and yellow) (New Horizons Entomology Services, Upper Marlboro, MD USA). Four-foot metal T-posts with metal wire were utilized to secure the bowls in place for sampling. Soapy water solution (water + a few drops of dish soap) was added to each bee bowl and bee bowls were set on two dates each month between 09:00 am and 011:00 am CST and the contents of bee bowls were collected after 24 hours. All bees were placed into 4oz Whirl Pak® (Nasco Fort Atkinson, WI) bags or glass vials containing 70% ethanol. In the laboratory, bees were identified to the level of genus using available taxonomic keys available online (Discover Life http://www.discoverlife.org and Bug Guide http://bugguide.net) and texts related to identification of native bees (Michener et al. 1994; Michener 2007; Wilson and Carril 2015). Bee species identification was conducted by Dr. Karen Wright (Texas A&M University). Voucher specimens of bees are currently held in part with Dr. Karen Wright (TAMU) and in the laboratory of Amede Rubio Texas A&M International University (Dept of Biology and Chemistry).

Ambient outdoor air temperature was collected monthly using a Kestrel 5000 Environmental Meter with LINK (Kestrel Meters, Boothwyn, PA) at each bee bowl cluster (n = 24) and averaged across samples to yield one value per sampling date. Annual and monthly accumulated rainfall data was accessed and downloaded from the NOAA weather database (https://water.weather.gov/precip/). Mean monthly temperature and rainfall was calculated and compared across years of sampling.

### Statistical Analyses

Data matrices (ecological habitat samples/plots x bee species) were constructed to calculate bee generic richness and bee total abundance (i.e. the total number of individual bees) to compare across habitats and time (i.e. two years). Counts of blooming plants were made monthly across the two-year study period. The row and column summary command in PCORD 7.0 (Wild Blueberry Media LLC, Corvallis, OR USA) was used to calculate diversity indices for both bees and plants. Shannon diversity (H) and Simpson diversity (D) was analyzed across habitats using one-way Analysis of Variance (ANOVA) (Magurran 1988) (JMP 14, SAS Institute Inc, Cary, NC).

Indicator species analysis (ISA) was used to determine indicator bee and plant taxa occurring in both habitat types. ISA compared the frequency and abundance of bees and plants among habitats to determine unique associations with statistical significance, which were calculated with permuted community data. Nonmetric multidimensional scaling (NMS) was utilized to ordinate the bee abundance data to determine associations between riparian and upland habitat types (n = 12 plots for each habitat) (PC-ORD, Corvallis, OR). Bees with less than three individuals were removed and abundance data was power transformed to the level of p = 0.5 square root prior to ordination. Sorensen distance measure was used, and 250 iterations were performed on real data. NMS analysis on abundance data suggested a two-dimensional solution following 51 iterations with a final stress of 18.48 and a final instability <10-5. Monte Carlo permutation show that the two-dimensional solution was significant (p=0.004). NMS and ISA were conducted using PCORD 7.0 (Wild Blueberry Media LLC, Corvallis, OR USA).

A mixed model with a repeated measures analysis was utilized (JMP 14, SAS Institute Inc, Cary, NC) to compare bee generic richness and total number of individuals among the riparian and upland habitats and over time. The model was constructed using the fixed main effects of bee generic richness and abundance, a full factorial between month and habitat, and a random effect of year with nested month (α = 0.05). Pooled data was used (across habitats) to determine relationships of bee generic richness and abundance, seasonality, and total bloom characteristics within the study area. Correlation analysis (i.e. non-parametric correlation Spearman’s ρ) was used to determine bivariate relationships among bee generic richness, bee abundance, blooming plant counts, and average temperature accumulated monthly precipitation (JMP 14, SAS Institute Inc, Cary, NC).

## Results

### Bee and Plant Community Summary

A total of 1,436 individuals representing 4 bee families, 28 genera and 68 species were collected across the riparian and upland habitats (Table 1). Halictidae (755 individuals) and Apidae (567 individuals) were the most speciose families with 25 species, followed by Andrenidae 10 species (74 individuals) and Megachilidae 8 species (40 individuals). The distribution of genera encountered were as followed: 15 (Apidae), 5 (Halictidae), and 4 (Andrenidae and Megachilidae). The 10 most dominant bee genera comprised 92 percent of the total number of individuals collected across the entire study, with 14 bee genera comprising the remainder of the community (8 percent of the total number of individuals of the community across two years). *Lasioglossum* had the highest number of individuals with 191 and was almost three times more abundant than *Apis* (68).

**Table 1.** Bee species summary of diversity indices in riparian and upland habitats.

| Species | Riparian | Upland |
| --- | --- | --- |
| *Agapostemon angelicus* | 37 | 38 |
| *Agapostemon melliventris* | 9 | 4 |
| *Ancyloscelis apiformis* | 8 | 16 |
| *Ancyloscelis sejunctus* |  | 1 |
| *Andrena macoupinensis* |  | 1 |
| *Andrena primulifrons* | 11 | 18 |
| *Andrena sp.A* |  | 23 |
| *Andrena sp.B* | 1 |  |
| *Andrena sp.D* |  | 1 |
| *Andrena sp.E* |  | 1 |
| *Anthophora californica* | 1 | 2 |
| *Anthophora occidentalis* | 21 |  |
| *Anthophorula compactula* | 3 | 2 |
| *Apis mellifera* | 141 | 72 |
| *Ashmeadiella cactorum* |  | 5 |
| *Ashmeadiella maxima* |  | 3 |
| *Ashmeadiella meliloti* | 1 | 9 |
| *Augochlorella aurata* | 4 | 2 |
| *Augochloropsis metallica* | 11 | 5 |
| *Calliopsis subalpina* | 9 | 3 |
| *Centris atripes* |  | 2 |
| *Ceratina shinneri* | 31 | 14 |
| *Diadasia diminuta* | 74 | 18 |
| *Diadasia enavata* | 1 | 1 |
| *Diadasia ochracea* |  | 2 |
| *Diadasia piercei* |  | 1 |
| *Diadasia rinconis* | 13 | 19 |
| *Epeolus sp.* | 1 |  |
| *Eucera lepida* | 3 | 1 |
| *Exomalopsis birkmanni* |  | 1 |
| *Florilegus condignus* | 2 | 1 |
| *Halictus ligatus* |  | 11 |
| *Halictus tripartitus* | 1 |  |
| *Lasioglossum (Dialictus) sp.K* |  | 1 |
| *Lasioglossum (Dialictus) hudsoniellum* | 4 | 18 |
| *Lasioglossum (Dialictus) nr. coactum* | 168 | 151 |
| *Lasioglossum (Dialictus) semicaeruleum* | 15 | 26 |
| *Lasioglossum (Dialictus) sp.* | 1 |  |
| *Lasioglossum (Dialictus) sp.A* | 10 | 13 |
| *Lasioglossum (Dialictus) sp.B* |  | 6 |
| *Lasioglossum (Dialictus) sp.C* | 1 |  |
| *Lasioglossum (Dialictus) sp.D* | 29 | 63 |

**Table 1. Continued**
|  |  |  |
| --- | --- | --- |
| *Lasioglossum (Dialictus) sp.E* | 3 | 1 |
| *Lasioglossum (Dialictus) sp.F* | 1 | 2 |
| *Lasioglossum (Dialictus) sp.G* | 4 | 3 |
| *Lasioglossum (Dialictus) sp.H* |  | 1 |
| *Lasioglossum (Dialictus) sp.I* | 47 | 31 |
| *Lasioglossum (Dialictus) sp.J* | 1 | 1 |
| *Lasioglossum (Dialictus) sp.K* | 5 | 3 |
| *Lasioglossum morrilli* |  | 1 |
| *Lasioglossum sp.L* | 16 |  |
| *Lasioglossum (Evylaeus) sp.* | 3 | 4 |
| *Lithurgopsis gibbosa* | 1 |  |
| *Lithurgopsis littoralis* | 4 | 1 |
| *Megachile brevis* |  | 1 |
| *Megachile sidalcea* |  | 2 |
| *Melitoma sp.* | 15 | 17 |
| *Mellisodes communis* | 1 | 3 |
| *Mellisodes tepaneca* | 26 | 38 |
| *Mellisodes tristis* | 1 | 2 |
| *Osmia subfasciata* | 5 | 8 |
| *Perdita ignota* | 2 | 1 |
| *Perdita sp.A* | 1 | 1 |
| *Pseudopanurgus sp.* | 1 |  |
| *Svastra obliqua* | 2 | 2 |
| *Tetraloniella spissa* | 2 |  |
| *Xylocopa tabaniformis* | 2 | 2 |
| *Xylocopa varipuncta* | 2 |  |
| Total Richness | 50 | 57 |
| Shannon | 2.80 | 3.03 |
| Simpson | 0.89 | 0.92 |

Across the total sampled area (upland and riparian habitats), a total of 57 flowering plants species with blooms were counted representing 24 families (Table 2). Fifty flowering plant species were counted in the riparian and twenty-four in the upland habitat. Sunflower (*Helianthus annuus*), narrowleaf globemallow (*Sphaeralcea angustifolia*) and silverleaf nightshade (*Solanum elaeagnifolium* Cavanilles) (Solanales: Solanaceae) were the dominant blooming plants observed in the study. Bee abundance and vegetation data were summarized graphically (Fig. 2)

**Fig. 2.**
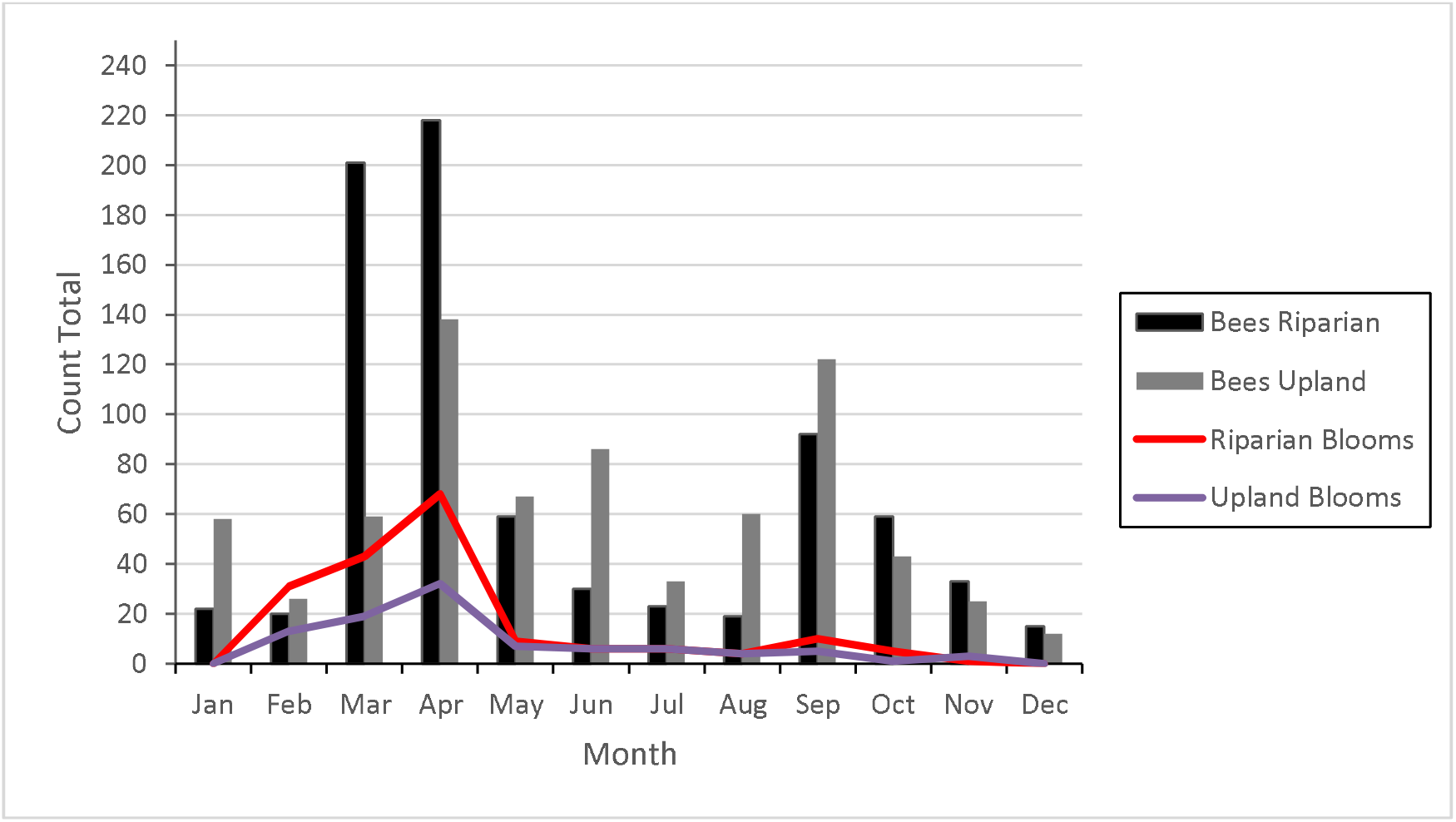
Comparison of total bee abundances and total bloom counts in upland and riparian habitats during the study period. Data has been pooled by years and shown as totals by month. Generic bee abundances were significantly correlated to total bloom counts (r_s_= 0.6260; *p* = 0.0006) (α = 0.05).

**Table 2.**
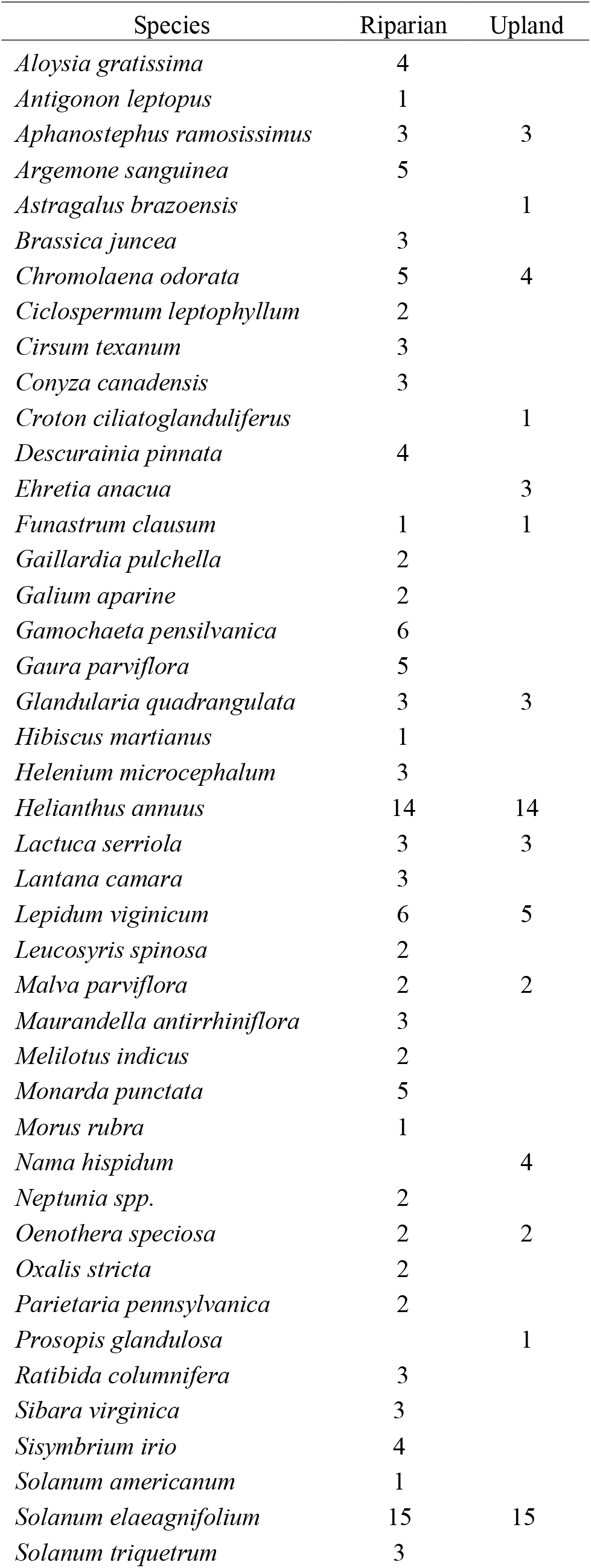
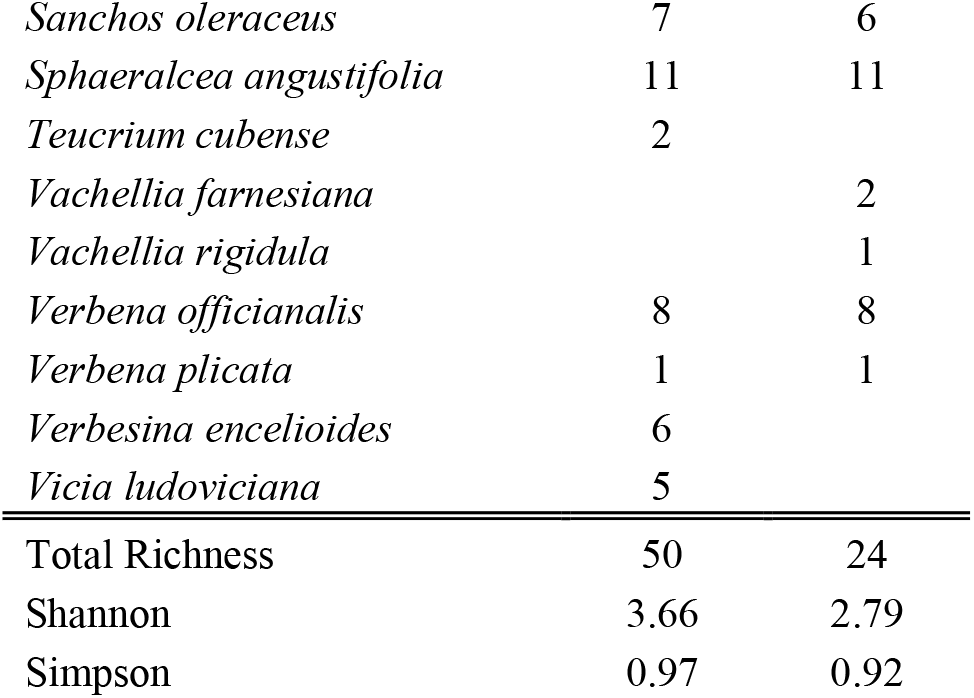
Blooming plant count and summary of diversity indices in riparian and upland habitats.

### Habitat Level Community Analyses

The top two dominant bee genera regarding abundances *Lasioglossum* and *Apis* accounted for 59 percent of the total number of individual bees collected. Pooled riparian and upland community data showed that bee generic richness between both habitats was comparable. Calculated Shannon and Simpson diversity between habitat types were very similar: bee Shannon diversity in riparian (2.80) and upland (3.03) and Simpson diversity in riparian was (0.89) and upland (0.92) (Table 1). ANOVA of Shannon (F = 3.56; df = 1, 22; *P* = 0.0723) and Simpson diversity (F = 2.70; df = 1, 22; *P* = 0.1145) were not significantly different between riparian and upland habitats. Three bee species were found to be significant indicator taxa; two in the riparian habitat *Anthophora occidentalis* (IV = 58.3; *P* = 0.0056); *Lasioglossum sp.L* (IV = 41.7; *P* = 0.034) and one showing affinity for the upland habitat *Halictus ligatus* (IV = 66.7; *P* = 0.0018).

Among 57 flowering plant species recorded, 7 were only encountered in the upland and 35 were unique to riparian habitats. A total of six plant species were most frequently encountered occurring in over 50 percent of sample plots and blooms of these species persisted an average of four months across all years in the current study: silverleaf nightshade (*Solanum elaeagnifolium*) (April-September), common sunflower (*Helianthus annuus*)(April-September), *Sphaeralcea angustifolia* (narrowleaf globemallow) (March-July), annual sowthistle (*Sonchus oleraceus*) (February-April), Texas vervain (*Verbena officinalis* ssp. *halei* Small) (Lamiales: Verbenaceae) (February-April), and cowpen daisy (*Verbesina encelioides* Cavanilles) (Asterales: Asteraceae) (March-May). Indicator species analysis of vegetation showed that three species were significant indicator species in riparian habitats, spiny pricklepoppy (*Argemone sanguinea*) (IV = 20; *P* = 0.046), spotted beebalm (*Monarda punctata*) (IV = 20; *P* = 0.048), and Pennsylvania cudweed (*Gamochaeta pensylvanica*) (IV = 20; *P* = 0.05). Calculated Shannon diversity of blooming plant presence was higher in riparian (3.66) than upland (2.79) plant communities (Table 2). However, subsequent ANOVA on Shannon diversity (F = 3.72; df = 1, 44; *P* = 0.0602) and Simpson diversity (F = 2.12; df = 1, 44; *P* = 0.15) showed differences in plant communities were not significant.

Habitat plots and bee abundances were ordinated using 2-dimensional NMS (*I* = 0.562 and *A* = 0.272). Ordination produced a significant result (Monte Carlo p = 0.004), with habitat plots distributed throughout the ordination space (Fig. 3). The horizontal axis (Axis 1) accounted for 51% and vertical axis (Axis 2) 78% of variance in the distance matrix.

**Fig. 3.**
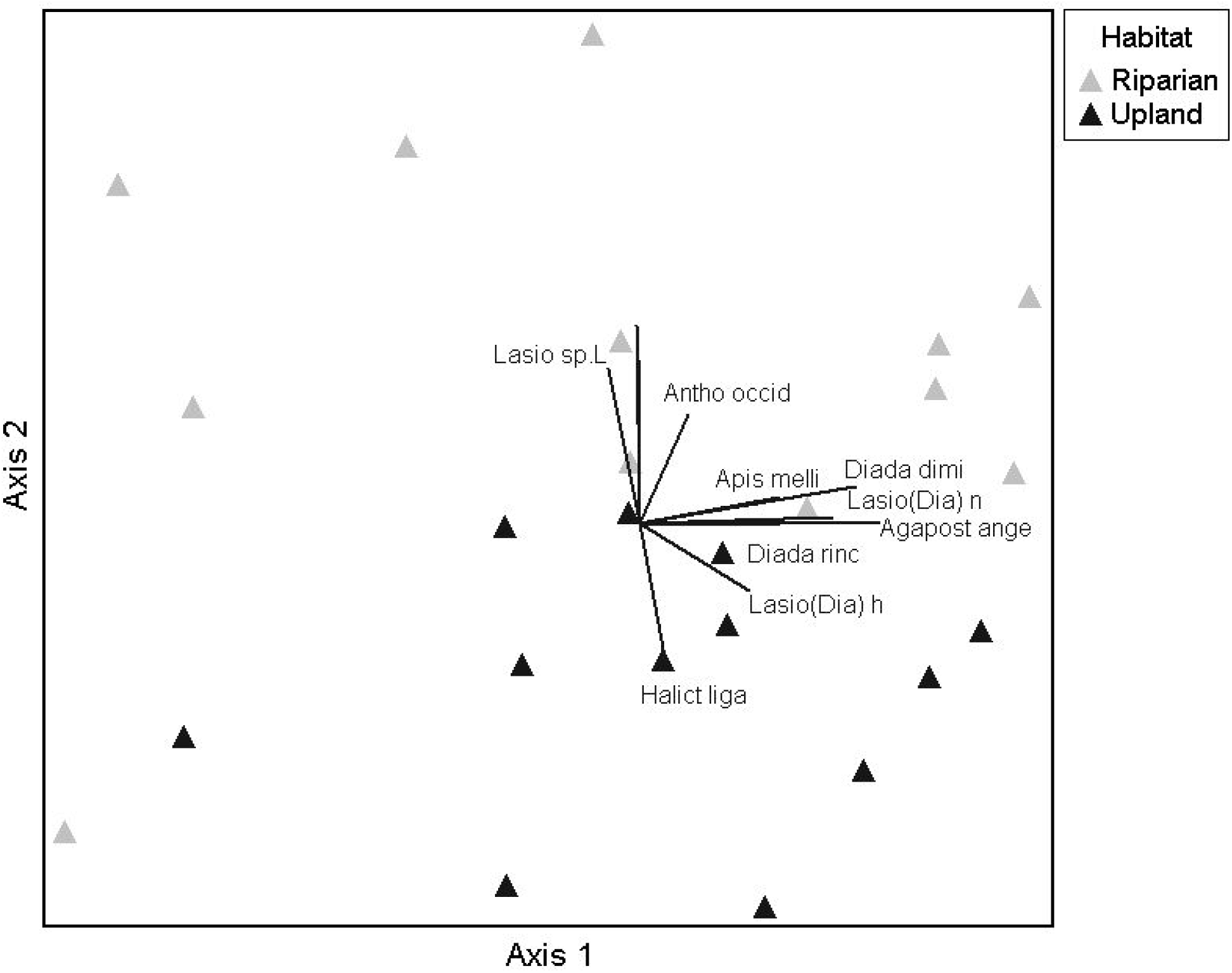
NMS ordination of bee abundance in riparian and upland habitat plots. Ordination is showing the affinity of *Lasioglossum sp. L* and *Anthophora occidentalis* for riparian habitats and *Halictus ligatus* for upland habitat type.

### Seasonal Bee Communities

Analysis of pooled bee abundance data showed no significant difference in abundance between years of data collection (F = 0.22; df = 2,22; *P* = 0.8015) (α = 0.05). However, effect tests in our statistical model showed a significant difference for monthly abundance, across all sites and years (F = 4.91; df = 11,12; *P* = 0.0048) (α = 0.05). Pooled monthly community data showed three peaks of higher bee abundance in the months of March (*P* = 0.0170), April (*P* = 0.0001) and September (*P* = 0.0139) (α = 0.05) (Fig. 4A). The two most abundant bees showed clear peaks, with *Apis* had the highest peak abundances in March and April and *Lasioglossum* in September.

**Fig. 4.**
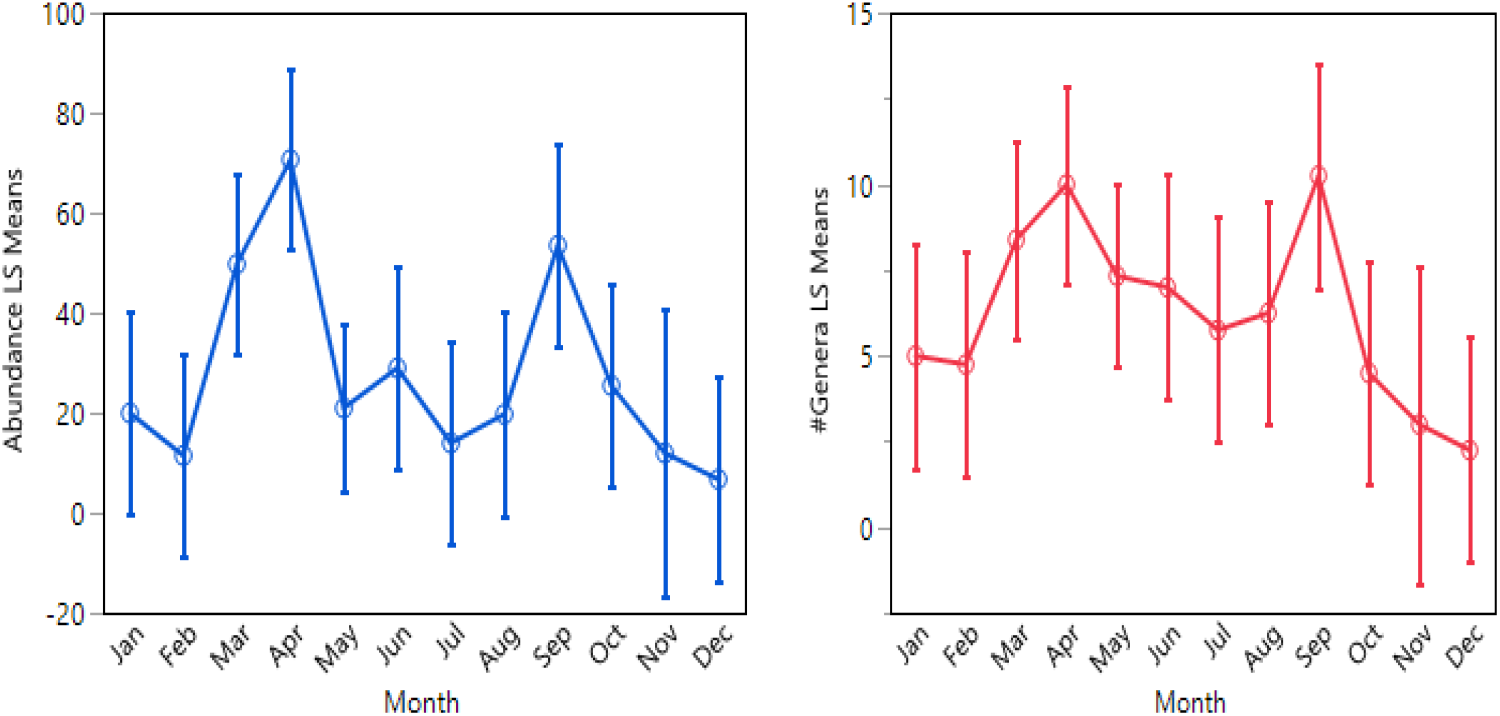
Interaction plots of pooled bee abundance by month (years combined) (A), and interaction plot of pooled generic richness by month (years combined) (B). Error bars represent 95% confidence intervals.

Analysis of pooled community data showed no significant difference in generic richness between years of data collection (F = 1.59; df = 2,21; *P* = 0.2262) (α = 0.05). Effect tests in our statistical model showed a significant difference for monthly generic richness, across all sites and years (F = 2.83; df = 11,11; *P* = 0.0473) (α = 0.05). Pooled monthly community data by month showed a bimodal trend of increasing generic richness in the months of April (*P* = 0.0115) (averaged 17 genera) and September (averaged 20 genera) (*P* = 0.0162) (α = 0.05) (Fig. 4B).

### Correlations of Bee, Blooming Plant Counts, Temperature and Precipitation

Riparian and Upland plant, bee, temperature, and precipitation data were pooled across years prior to analysis. There was a strong positive correlation between bee generic richness and blooming plant counts, which were statistically significant, (r_s_ = 0.6057; *P* = 0.0010) (α = 0.05) (Fig. 5A). Similarly, bee abundance was positively correlated with blooming plant counts and statistically significant, (r_s_= 0.6298; *P* = 0.0006) (α = 0.05) (Fig. 5B). Average monthly temperature was positively correlated with bee generic richness and statistically significant (r_s_ = 0.4566; *P* = 0.0190) (α = 0.05) (Fig. 5C). Conversely, bee abundance (r_s_ = 0.0971; *P* = 0.6370) (α = 0.05) and blooming plant counts (r_s_ = 0.1161; *P* = 0.5722) (α = 0.05) were not correlated with temperature. Accumulated monthly precipitation (cm) was positively correlated with bee abundance and statistically significant (r_s_ = 0.4005; *P* = 0.0426) (α = 0.05) (Fig. 5D). However, accumulated monthly precipitation (cm) was not positively correlated with bee generic richness (r_s_ = 0.2806; *P* = 0.1650) (α = 0. 05) and blooming plant counts (r_s_ = 0.1658; *P* = 0.4184) (α = 0.05).

**Fig. 5.**
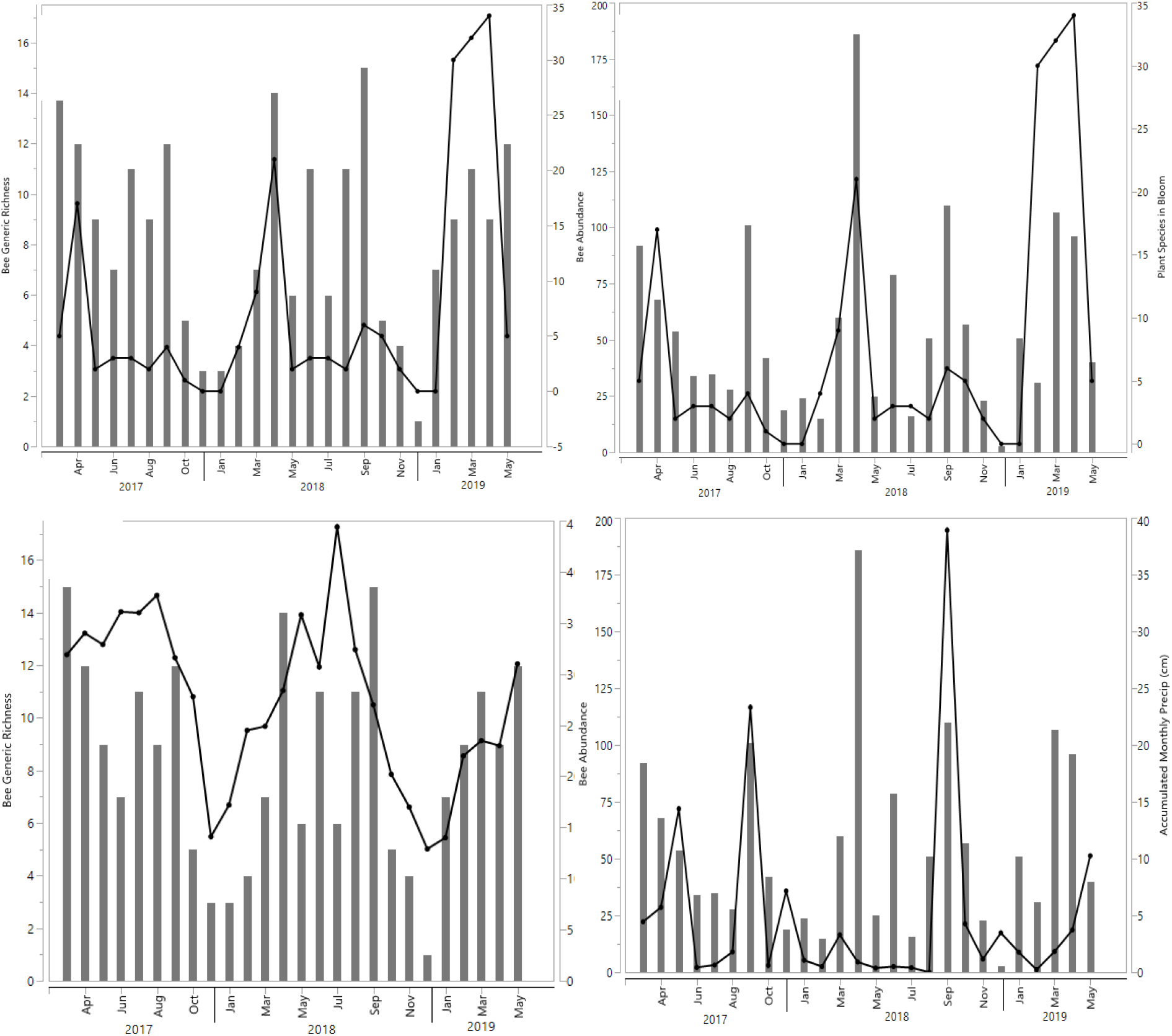
Overlay plot of blooming plant counts (line) and bee generic richness (bars) (A), blooming plant counts (line) and bee abundance (bars) (B). Overlay plot of monthly average temperature (line) and bee generic richness (bars) (C) accumulated monthly precipitation and bee abundance (bars) (D).

## Discussion

The goal of community sampling was to census the seasonal abundances and diversity of bees using the riparian corridor for foraging or other behaviors such as nesting preference (Fellendorf, Mohra, and Paxton 2004). Our study shows that diverse native bees are utilizing riparian habitat resources and if present trend of anthropogenic disturbances continue, this could have a significant impact on extant bee communities. To date, the current study provides the only account of bee diversity and flowering plant diversity for this important habitat in the region.

Bee data may have been biased towards some families such as Halictidae because of bee bowl sampling (Hall 2016). However, this family is commonly abundant and represents a large portion of native biodiversity in the region that could benefit from warm, sandy soils and diverse flowering vegetation (Michener 2007). Among the dominant genera of bees collected in the study, *Lasioglossum* was most abundant. They are ground nesting bees and can have an array of social behaviors that range from strictly solitary to parasitic (Michener 2007). The high number of collected *Lasioglossum* may have also been attributed to bee bowl sampling (Roulston, Smith, and Brewster 2007). Bees in the genus *Diadasia* (Patton) (Hymenoptera: Apidae) are small to large sized hairy bees that range in size from 5-20 mm (Michener 2007). The bees encountered from this genus were observed mainly foraging on narrowleaf globemallow (*Sphaeralcea angustifolia*) which was a common plant present in both habitat types. Many of the bees in this genus are foraging specialists and make shallow nests often with tubular entrances around the opening (Michener 2007). The genus *Melissodes* (Latreille) (Hymenoptera: Apidae) are medium to large bodied bees 7.5-16 mm (Michener 2007). Many of the *Melisodes* collected were in early - mid fall (September-October) which is a commonly characteristic of this genus (Wilson and Carril 2015). The bees in this genus are specialists that primarily forage on flowers of the family Asteraceae, but few may be generalists (Michener 2007). All *Melissodes* are ground nesting solitary bees (Michener 2007). Unexpectedly, September showed are large spike bee generic richness and abundance although blooming plants remained low. Upon further investigation, blooming invasive plant San Miguelito vine (*Antigonon leptopus* Hooker and Arnott) (Polygonales: Polygonaceae) was found growing within the riparian habitat along with other dominant flowering plant species. The availability of blooms by common sunflower, silverleaf nightshade and presence of San Miguelito vine may provide resources for late season bees such as *Melissodes*.

The NMS ordination showed a clear presence of heterogeneity among bee species utilizing upland and riparian habitats. Habitat heterogeneity was further supported by the results of the indicator species analysis where three bee species were found to be significant indicator taxa. *Anthophora occidentalis* and *Lasioglossum sp.L*, showed indication for the riparian habitat and *Halictus ligatus* in the upland habitat. Species of the genus *Anthophora* are robust, fast flying bees that exclusively nest in banks or flat ground (Michener 2007). *Anthophora occidentalis* may also have benefited from a nearby water source like the Rio Grande since it is known regurgitate water to moisten soil during excavation of the nest. Species *Halictus ligatus* is part of a large genus of bees that is very diverse and like many other native bees in this group are ground nesting. This species may have had a strong indication for upland habitat due to the higher presence of bare ground caused by the aggregate growth habit of invasive buffelgrass (*Cenchrus ciliaris*). *Halictus ligatus* may have preferred the flat, compacted soil in these sites as a suitable nesting habitat.

Riparian habitats recorded two times more flowering plant species than upland habitats, which likely stimulated upland bees to forage in the riparian zone. This is further supported by distances between the two zones which averaged only 172 m, which might not have been enough spatial distance to present differences in bee capture. Consequently, the proximity of both habitats created overlap of similar plant communities in which would be within bee foraging range. In a study conducted by Gathmann and Tscharntke (2002) showed that bees averaged 150 – 600 m of foraging distance between nesting sites and floral resources, which comparatively is well within our measured distance between habitats. Other covariables that drive distances between habitats, elevation, and distance to river, likely in part drive soil and plant differences in riparian and upland habitats.

Overall, results show high similarities among habitats and dominant, soil-nesting bees in both habitats. The succession of invasive grasses, primarily giant reed, is a dynamic process driven by disturbance. As giant reed continues to spread and create large monotypic stands, floral diversity and potential pollinator/bee resources may decline (Herrera and Dudley 2003). This may cause extirpation of rare species from the riparian corridor. Furthermore, investigating how disturbances affect soil nesting native bees would advance our understanding of bee biology in a unique riparian community. Further, ecological restoration involving native plants could assist in management of invasive giant reed, coupled with other benefits from intensified riparian management involving giant reed (Patiño et al. 2018). Ecological restoration towards native vegetation could support initiative for management of the corridor by reestablishing native vegetation to replace dense stands of giant reed.

Seasonally, bee communities can vary significantly over time, largely depending on the availability of floral resources, seasonal phenology, and environmental factors (Kimoto et al. 2012). Bee genera and abundance showed a high seasonal/monthly variation but conversely not significant inter-annual differences. Lack of significant inter-annual differences in bee diversity may be in part due to the regions relatively consistent subtropical climate, which in turn may develop patterns in bee behavior (Boucek et al. 2016). Generic richness across months and years, were significantly different especially in April and September. Similarly, abundance across months and years was significantly different with March, April and September having the highest bee abundance which may be largely attributed to floral availability and temperature (Classen et al. 2015). Kimoto et al. (2012) showed similar trends in their study where during the spring growing season had the highest bee activity which was also strongly associated with available floral resources and average monthly temperature. In our study, temperature extremes may have negatively affected bee behavior, since the data showed decreased abundance and generic richness at temperatures below 15°C and above 30°C. To support this, blooming plants and bee generic richness are strongly correlated in the months of April and September which may indicate an optimal temperature range of 25°C −30°C (Fig. 5C). Temperature extremes could have limited bee access to floral resources although they were abundant. Bee abundance increased with accumulated monthly precipitation in the month of September (across years) (Fig. 5D), but unexpectedly was not associated with other study variables like genera richness and blooming plant counts. September rainfall may have initiated blooming response in dominant plants that may have caused increased foraging in seasonal bee genera.

Along a narrow two mile stretch of the LRG we recorded previously undocumented bee and flowering plant communities, which supports further studies and conservation actions involving this important river and its riparian corridor. How wild and native bees use this habitat remains an important area of investigation, especially considering intensified management in the riparian corridor. The community approach and findings of the current study show diverse bees using resources provided in this variable habitat, while the diversity and areal coverage of flowering plant communities in the riparian are likely affected by competition from highly invasive plants such as giant reed. These environmental flow-mediated habitats are facing additional severe threats from anthropogenic activity and invasive plant species (Fowler et al. 2018). The flowering plant communities, soil structure (affecting bee nesting) and bee communities could serve as biological targets for ecological restoration conducted in this intensively managed riparian corridor.

